# Exposure to *Pseudomonas spp.* increases *Anopheles gambiae* insecticide resistance in a host-dependent manner

**DOI:** 10.1101/2023.11.13.565999

**Authors:** Luís M. Silva, Gwendoline Acerbi, Marine Amann, Jacob C. Koella

## Abstract

The microbiota of mosquitoes influences many aspects of their biology, including developmental processes, mating and sexual reproduction, immune functions, and refractoriness to pathogens. Here, we considered their role in resistance against insecticides. In particular, we assessed how larval infection of a permethrin-resistant and a sensitive colony of *Anopheles gambiae* by four strains belonging to three different *Pseudomonas* species affects several life history traits and the impact of the insecticide on adult mortality. Our data showed that all four *Pseudomonas* strains persisted in adults until death. The bacteria increased the likelihood that mosquitoes survived 24 hours after exposure to permethrin by up to two-fold. The impact of the bacteria depended on the bacterial strains and the mosquito colony: in the resistant colony, all bacteria increased survival by about 2-fold, while in the sensitive colony, only two of the four strains increased survival. The benefit concerning insecticide resistance came with little to no impact on the other traits (i.e., larval mortality, developmental time and adult longevity). Altogether, our results highlight the importance of considering environmental microbial exposure and mosquito microbial communities in epidemiological and vector-control studies, while also suggesting a possible role for *Pseudomonas spp.* as a symbiont in *A. gambiae*.

## Introduction

Although the response of mosquitoes and other insects to insecticides has a clear genetic component^1^, it is also influenced by the environment. Thus, adult mosquitoes’ likelihood to be killed by insecticides has been shown to be affected by temperature at exposure^2^ or diet^3^. An important component of the environment of mosquito larvae is the community of bacteria that occur in water, sometimes at high concentrations^4–9^. These bacteria are often ingested by mosquito larvae. While some of them are digested and serve as food, others can persist and may contribute to the microbiota of mosquitoes. Since microbiota often survives pupal development, these aquatic bacteria can affect the composition of the microbiota in adults^10–12^, which in turn affects life history traits^7,13–16^ and other traits, such as insecticide resistance^13,16–18^. It should be noted, however, that most studies are correlational, associating the microbiota composition found in a mosquito to its performance, rather than experimental^13,16,17^. An exception is the bacterium *Asaia*, commonly found in mosquito microbiota, whose insecticide-degrading mechanisms have been studied in controlled experimental settings^18^. The paucity of experimental data biases our predictions and makes it difficult to quantify the role of the microbiota in insecticide resistance.

One of the most common groups of bacteria in the microbiota of mosquitoes is *Pseudomonaceae*, which includes the ubiquitous genus *Pseudomonas*^8,19,20^. The high frequency of these bacteria may reflect that *Pseudomonas* gives mosquitoes a particular benefit. Indeed, at least some species can degrade certain insecticides^21–25^. However, the impact of these bacteria on insecticide resistance (or other life history traits) of mosquitoes has been largely overlooked. We, therefore, decided to test whether *Pseudomonas* species affects the insecticide resistance or important life history traits of two colonies of the mosquito *Anopheles gambiae* (the insecticide-sensitive Kisumu colony and permethrin-resistant RSP colony). We used four members of the *Pseudomonas* genus: *P. aeruginosa*, *P. entomophila* and two strains of *P. fluorescens. P. aeruginosa* is a pathogen not only of insects but also of humans, with a case-fatality rate of 40%, partly due to multi-resistance to antibiotics^26^. *P. entomophila* is particularly toxic to insects^27,28^, as it produces a toxin called Monalysin^29^, which forms pores in the peritrophic matrix of the gut, enabling bacterial cells to invade the rest of the body. *P. fluorescens* has particular relevance for disease control in plants^30^ and it is less virulent in insects than the other two species^31^.

All of the bacterial strains were isolated from environmental sources, such as soil. As an insecticide, we selected permethrin, which is a widely used pyrethroid in vector control. Using a permethrin-resistant and sensitive colony of mosquitoes, we aimed to test: i) if environmental exposure to these bacteria in a laboratory setting leads to infection, ii) if the infection persists within the mosquitoes, iii) if the mosquitoes’ life histories are affected by the exposure, infection, or persistence and, most importantly, iv) whether the bacteria affect insecticide resistance.

## Results

*Developmental traits*: Infection by bacteria increased larval mortality of RSP mosquitoes slightly, with up to 5% of the larvae dying (χ2 = 12.796, df = 4, *p* = 0.012), but no other statistically significant differences in developmental traits were observed in either mosquito colony (Fig. 1). In particular, longevity was not affected by bacterial exposure for either strain of mosquitoes (Kisumu: *df* = 4, χ2 = 3.281, *p* = 0.512; RSP: *df* = 4, χ2 = 3.049, *p* = 0.550; Fig. 3).

**Figure 1.**
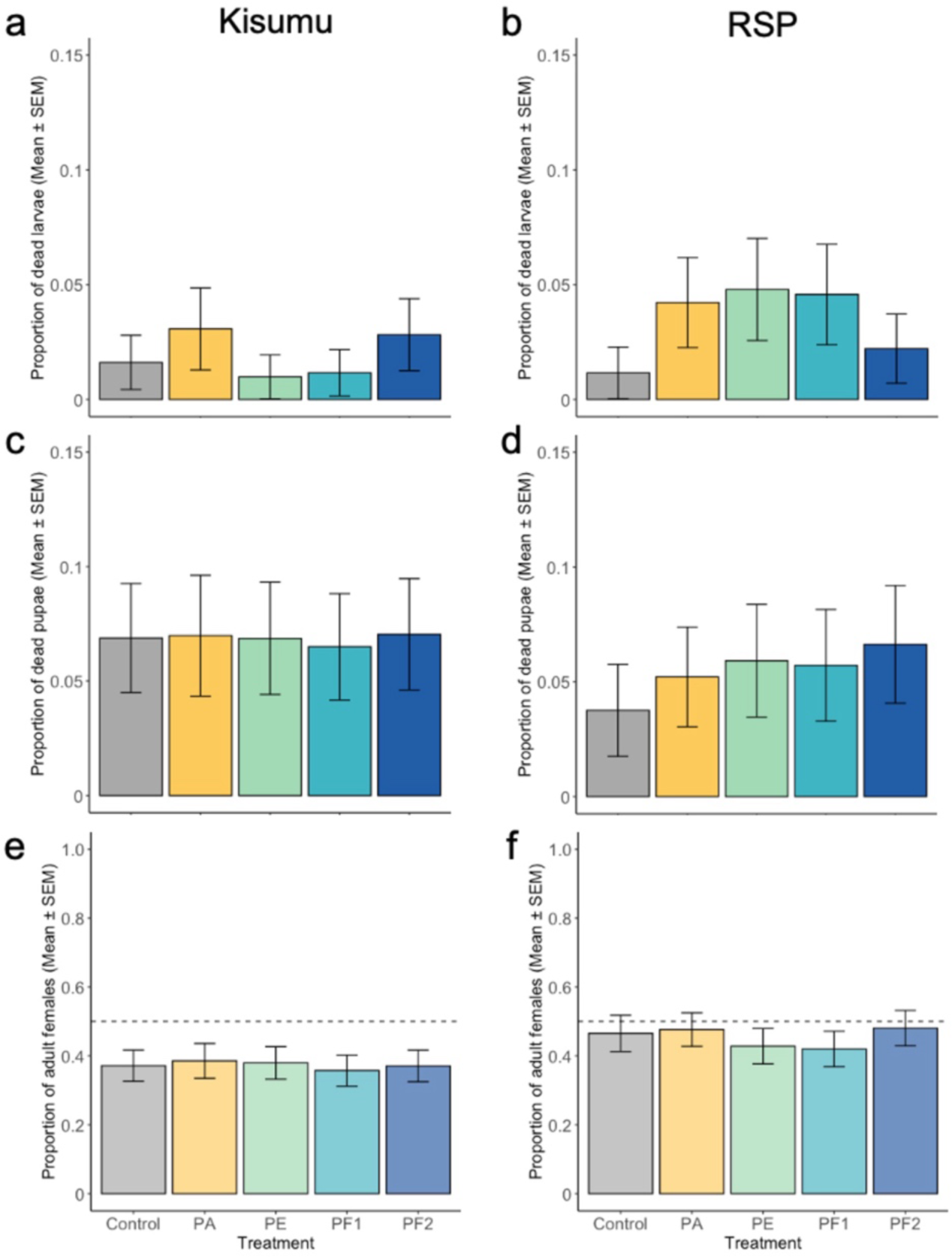
Developmental traits. **(ab)** Larvae mortality, as the proportion of dead larvae. Sample size is as follows: Control 436 and 346, PA with 358 and 403, PE with 408 and 355, PF1 with 431 and 350, PF2 with 326 and 362 individuals for Kisumu and RSP colonies, respectively. **(cd)** Pupae mortality, as the proportion of dead pupae. Sample size as follows: Control with 429 and 342, PA with 347 and 386, PE with 404 and 338, PF1 with 426 and 334, PF2 with 414 and 354 individuals for Kisumu and RSP colonies, respectively. **(ef)** Sex-ratio, as the proportion of females. The sample size is as follows: Control with 399 and 329, PA with 322 and 365, PE with 376 and 317, PF1 with 398 and 314, PF2 with 384 and 330 individuals for Kisumu and RSP colonies, respectively. The dashed line at 0.5 signifies an equal number of males and females. See Supplementary Table S1 for further details on the statistical analyses.

*Persistence of infection*: In mosquitoes that survived up to the test, the bacteria load was similar for the four bacteria strains (*df* = 3, F = 0.912, *p* = 0.437) and did not decrease from three days after emergence to ten days (*df* = 1, F = 1.107, *p* = 0.295) (Fig. 2a). In dead mosquitoes of the RSP colony, bacterial load was also similar for the four bacteria (*df* = 3, F = 0.535, *p* = 0.662) (Fig. 2c), but in the sensitive Kisumu colony the load among bacteria strains (*df* = 3, F = 25.903, *p* < 0.001), with PA having the highest load and PE2 the lowest (Fig. 2b). Note that only 39 RSP mosquitoes died, making it difficult to reach strong inferences. For further details on the statistical analysis and post-hoc tests see Supplementary Table S2.

**Figure 2.**
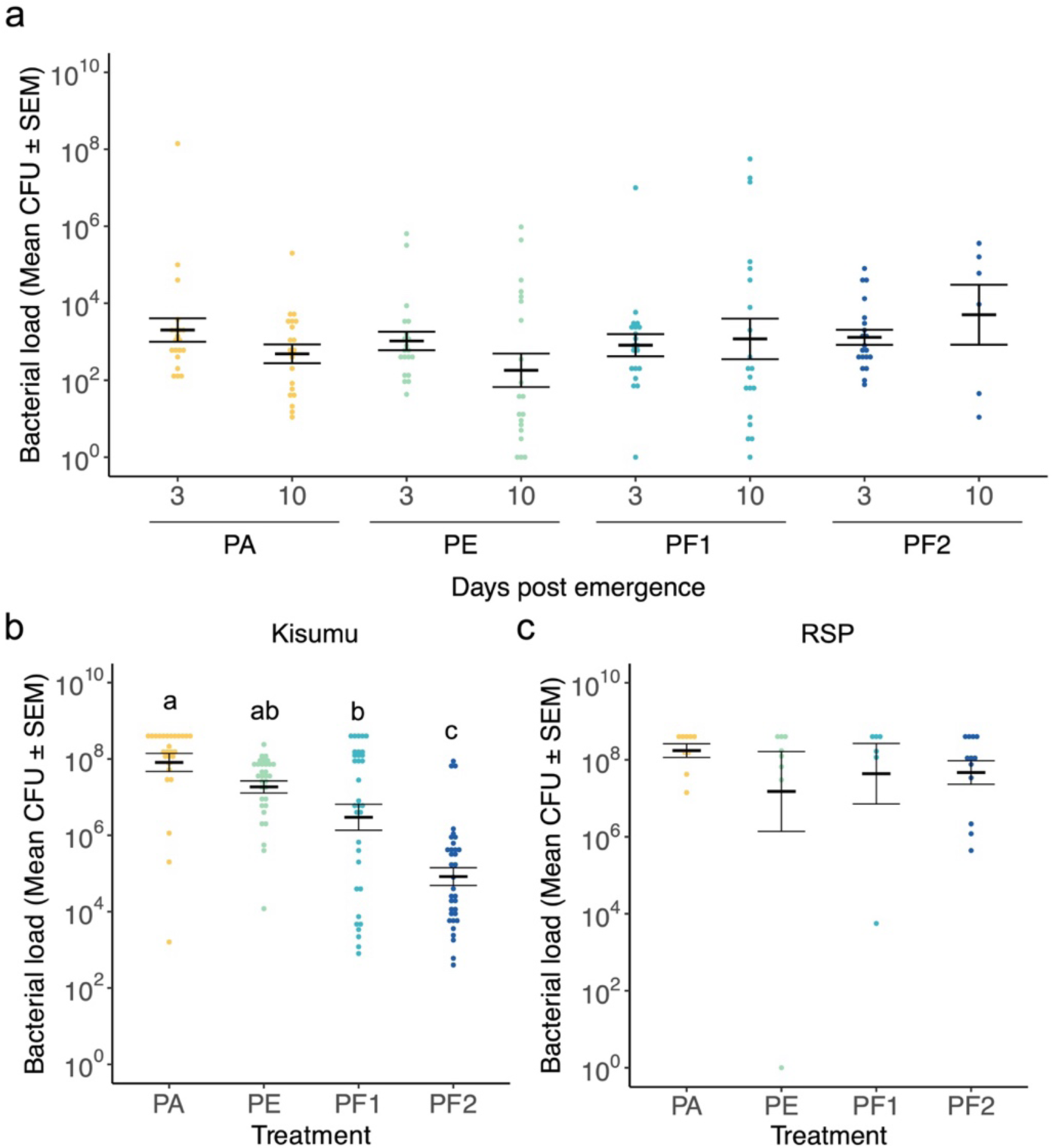
Bacterial persistence. **(a)** Individual bacterial load per surviving female mosquito after exposure to one of the four bacterial strains three days or ten days after emergence. Each treatment and time point included 20 females except for PF2 at day 10, for which we could test only six females. **(bc)** Individual bacterial burden for mosquitoes that died between 20 and 30 days after emergence for Kisumu and RSP. The sample sizes were: PA with 27 and 9, PE with 32 and 8, PF1 with 33 and 6, and PF2 with 34 and 12 females in Kisumu and RSP, respectively. Statistical analysis and multiple comparisons can be found in Supplementary Table S1. Different letters denote means that are significantly different from one another.

*Permethrin resistance*: In both mosquito colonies the bacterial infection tended to decrease the likelihood that mosquitoes died within 24 hours of exposure to permethrin. The exposure treatment affected both host colonies (Kisumu: *df* = 4, χ^2^ = 19.847, *p* < 0.001; RSP: *df* = 4, χ^2^ = 24.802, *p* < 0.001). A post-hoc analysis showed that in the Kisumu colony, PE survived better than the control (z-ratio: 2.720, *p* = 0.031), while in the RSP colony, PA (z-ratio: 3.054, *p* = 0.019), PE (z-ratio: 4.186, *p* < 0.001) and PF1 (z-ratio: 4.263, *p* < 0.001) survived better comparatively to the control. For further details on the statistical analysis and post-hoc tests see Supplementary Table S4.

## Discussion

Although several correlational studies have described an association between symbiotic microbes and insecticide resistance of mosquitoes^13,14,16,18^, experimental evidence for the link is lacking. We helped fill the experimental gap by showing that *A. gambiae* experimentally exposed as larvae to *Pseudomonas* strains are then infected throughout their lives and some of them provide resistance to permethrin with little cost concerning larval mortality or adult longevity.

While each of the bacterial strains infected, both, Kisumu and RSP mosquitoes up to their death, the bacterial load was influenced by the combination of the bacterial strain and the mosquito colony. In particular, it differed among bacterial strains in Kisumu but not in RSP mosquitoes. While our result corroborates a suggestion that the genotype of a bacterium’s host could drive differences in bacterial load upon death^32^, we note that we assayed only 20- to 30-day-old mosquitoes of adulthood and that only a few RSP mosquitoes survived up to this age.

Although the bacterial infection persisted throughout the mosquitoes’ life, it had no impact on longevity (Fig. 3), pupal mortality or sex ratio (Fig. 1). There was, however, a small (but insignificant in multiple comparison tests) effect of infection on larvae mortality in RSP, with mortality increasing from 1% in uninfected larvae to 5% for PE-infected larvae. This result is in striking contrast to the high mortality of *Drosophila melanogaster* ^27,28^ after septic and/or oral exposure to this genus. Our main result is that mosquitoes infected with *Pseudomonas* bacteria were better able to survive exposure to permethrin than uninfected mosquitoes. The bacteria had a stronger impact on the resistant RSP mosquitoes than the sensitive Kisumu ones. Thus, in RSP the difference between infected and sensitive mosquitoes was greater than in Kisumu, and three bacteria increased resistance in RSP, but only one did so in Kisumu. While this phenomenon has been observed before with other microbes of mosquitoes^18,33^, such as *Asaia*, this is the first study showing that *Pseudomonas* increases the insecticide resistance of *A. gambiae*. One possible mechanism is that the bacteria can degrade permethrin themselves. Pyrethroid degradation mechanisms have been described for *Asaia*, which can degrade them with pyrethroid hydrolase genes that are present in mosquito-symbiotic strains but not in non-symbiotic strains^34^. Some members of *Pseudomonas are* rich in hydrolases^35^, such as esterases, and several can degrade different insecticides in various hosts^21,23–25,36^. For example, a strain of *P. aeruginosa* isolated from the cockroach *Blatta orientalis* degrades *in* vitro 88.5% of endosulfan-based insecticides in under 10 days^37^. In addition, the bacteria might stimulate the mosquito’s detoxification cascade. For instance, the permethrin-resistance of RSP is due to a combination of knockdown resistance and cytochrome P450 activity, and exposure to permethrin increases the expression of esterases^38^ and antioxidant enzymes^39^. Stimulation of any of these detoxifying mechanisms by *Pseudomonas* strains would explain why we observe more strains contributing to increased insecticide resistance in RSP than in Kisumu (Fig. 4). It is important, though, to highlight the differences in recent evolutionary history of the two colonies. While Kisumu has been maintained at larger numbers in the laboratory, RSP is subjected to frequent bottlenecks as a consequence of its selection protocol. Therefore, Kisumu is probably genetically more variable than RSP, which would increase the stochasticity of the insecticide resistance response and hinder the detection of any differences due to bacterial strains.

**Figure 3.**
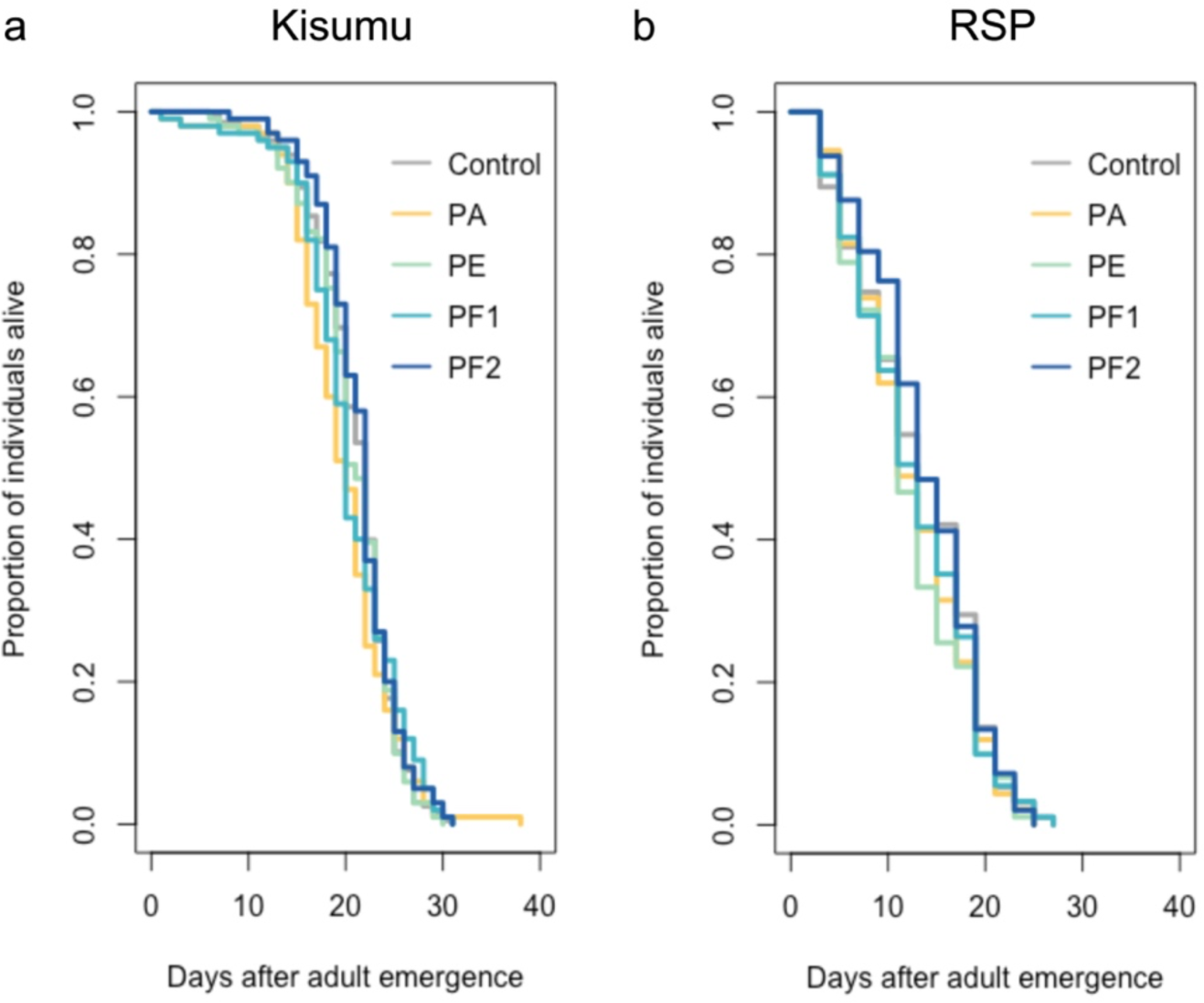
Adult longevity after exposure to Pseudomonas *spp*. Survival as a function of the day after exposure to one of four bacterial strains at the dose of 10^4^ CFU/ml. An uninfected control raised in distilled water was included. **(a)** Kisumu and **(b)** RSP mosquito colonies. The sample sizes are: control with 100 and 95, PA with 100 and 92, PE with 101 and 90, PF1 with 100 and 91, and PF2 with 100 and 97 females for Kisumu and RSP, respectively. Further details on the statistics are provided in Supplementary Table S2.

**Figure 4.**
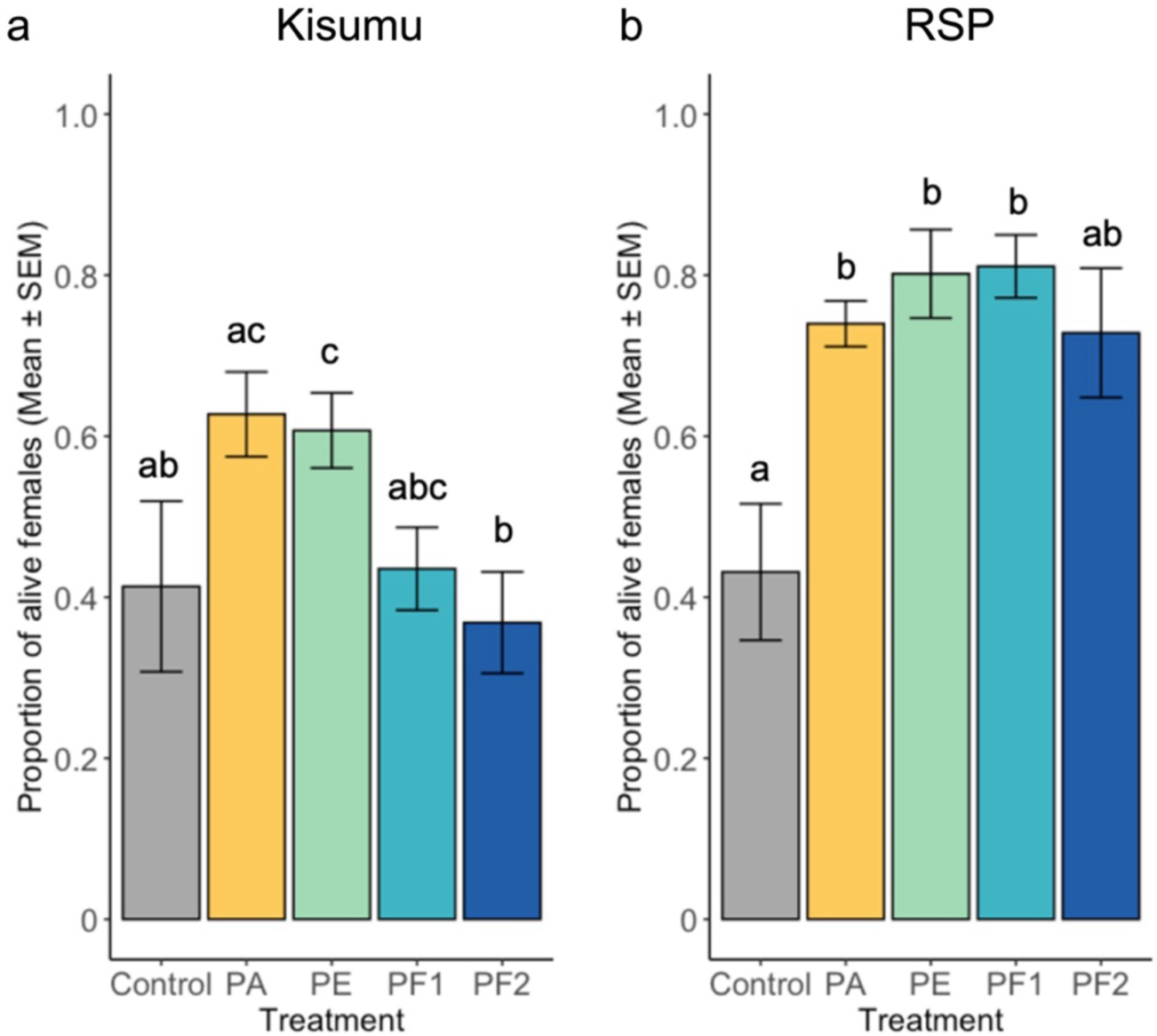
Insecticide resistance. Five-day-old individuals from the permethrin-susceptible colony Kisumu **(a)** and permethrin-resistant colony RSP **(b)** were exposed for 10 or 30 min to permethrin, respectively, and their mortality was assessed after 24 hours. Exposure was performed in five groups of approximately 15 females per treatment and colony. The sample sizes were: control with 73 and 68, PA with 81 and 82, PE with 86 and 101, PF1 with 85 and 92, and PF2 with 98 and 92 females for Kisumu and RSP, respectively. A multiple comparisons test was conducted to assess differences between treatments, which are denoted by different letters. Further details on the statistics are provided in Supplementary Table S3.

In summary, the high prevalence of *Pseudomonas spp.* in natural aquatic bodies combined with correlational and experimental evidence of an advantage against insecticides without a substantial cost with regard to life history traits in *A. gambiae* emphasizes the importance of considering the rearing environment and its consequences when studying and designing vector control strategies.

## Materials and Methods

### Experimental system

We used the permethrin-sensitive Kisumu colony^40^ and the permethrin-resistant RSP colony of *Anopheles gambiae (s.s.)*^38^. Resistance is due to a combination of knockdown resistance and increased activity of cytochrome P450. The colonies are maintained in our lab in three cages of overlapping generations (with an age difference of one week) with about 600 individuals per cage. To maintain permethrin resistance in RSP, L4 larvae are exposed to 0.5mg/L (Sigma-Aldrich, Inc., St. Louis, Missouri) for 24 hours every three generations. The experiment was started with eggs from each cage. The colonies and experimental individuals were maintained at 26.5 ± 0.5 °C and 70 ± 5% humidity, with a 12:12 light-to-dark photoperiod.

### Bacterial strains and infection assay

We used *P. aeruginosa* (Migula strain, DSM 50071), *P. entomophila* (DSM 28517), isolated from a wild-caught *Drosophila melanogaster*^41^, and two strains of *P. fluorescens* (Migula strain, DSM 50090 isolated from a pre-filter water tank; DSM 17397, isolated from tap water). Hereafter, we call the four strains: PA, PE, PF1 and PF2, respectively. PE was provided by Bruno Lemaitre (EPFL, Switzerland), while the others were provided by Saskia Bindschedler (UniNe, Switzerland). The bacteria were stored in 35% glycerol at -80°C, and new cultures were prepared for each experiment.

Bacterial preparation was adapted from Acuña-Hidalgo, Silva, Franz *et al.* ^27^. In brief, all bacterial species were plated in LB media (Luria/Miller broth, Carl Roth) for approximately 15h at 30°C, followed by an overnight liquid culture in LB media (Luria/Miller broth, Carl Roth) at a temperature of 30 °C and 200 rpm. After the overnight culture, the bacteria were washed twice by centrifuging the bacterial solution at 2880 × g at 4 °C for 10min, discarding the supernatant and replacing it by autoclaved distilled water. Their concentration was then estimated from the optical density (OD) of the bacterial solution measured with a Thermo Scientific BioMate 160 spectrophotometer at 600 nm. We added distilled water to the solution to obtain the desired concentration. To confirm the bacterial concentration, we plated and incubated the final bacterial solution and counted CFUs the following day.

Freshly hatched larvae (3-6 hours old) were reared in groups of 200 larvae in a glass tray (30 x 21 x 6 cm) containing 600 ml of one of the bacterial solutions or the negative control (distilled water). We gave the larvae Tetramin Baby ® fish food every day according to their age: 0.04 mg/larva on the day of hatching, 0.06 for one-day olds, 0.08 at age 2, 0.16 at age 3, 0.32 at age 4 and 0.6 at age 5 or older. We washed pupae twice in distilled water to remove residual bacteria from the body surface. This was performed by transferring sequentially the pupae with a disposable Pasteur pipette through two 500 ml beakers containing 400 ml of distilled water each, adapted from Silva (2017)^42^. We then put the pupae into 50 ml Falcon tubes, allowed them to emerge into adults, discarded the males, and moved females individually into plastic cups (5 cm diameter x 10 cm), where they stayed the remaining duration of their experiment and had constant access to a 6% sucrose solution.

### Bacterial load, development and longevity

In a preliminary experiment, we exposed the permethrin-sensitive colony (i.e., Kisumu) to 0, 10^2^, 10^4^ or 10^6^ CFU/ml of bacterial solution during larval development. These doses fall within the range of *Pseudomonas spp.* concentrations in aquatic bodies in which mosquitoes typically develop^43,44^. Three days after emerging, the mosquitoes were killed and the proportion of infected individuals was measured (Supplementary Fig. S1). Since many larvae died of infection by PE at a dose of 10^6^ CFU/ml, we limited the dose for all bacteria to 10^4^ CFU/ml in the following experiments. To test if any of the four bacterial strains can chronically infect *A. gambiae*, we thus exposed larvae to 0, 10^2^ or 10^4^ CFU/ml and measured the bacterial load three days (i.e., during the acute phase) and ten days (i.e., during the chronic phase) after emergence^32^. For each treatment, we measured bacterial load in 20 live mosquitoes at each time-point (Fig. 2a). Each mosquito was put into a 1.5 mL microcentrifuge tube containing 100 µl of distilled water and one stainless steel bead (Ø 5 mm). The mosquitoes were homogenised with a Qiagen TissueLyser LT at a frequency of 30 hz for one minute and then pelleted down for 10 seconds with a microcentrifuge. The resulting homogenate was diluted to 1:10^6^ of its original concentration and 50 µl of the following dilutions were plated in LB agar: 1:1, 1:10^2^, 1:10^4^, 1:10^6^. The CFU counts were done after incubating for 15 hours at 30 °C. For each mosquito we considered the CFU count from the lowest countable dilution to avoid additional dilution errors. The microbiota from our laboratory mosquitoes does not grow easily under these conditions. Nevertheless, as a control check, we also plated the uninfected individuals in all time points.

We then tested for differences in a set of developmental traits (i.e., larval and pupal mortality, as well as sex-ratio; Fig. 1) and longevity (Fig. 3) for permethrin-sensitive and -resistant colonies of mosquitoes in the presence or absence of each bacterial treatment. Individuals who died between day 20 and 30 after emergence were also plated, as described above, to confirm that they harboured bacteria when they died (Fig. 2bc).

### Exposure to permethrin

We assessed if any of the tested bacteria had an effect on the response to permethrin in permethrin-resistant or sensitive colonies. First, we tested if permethrin would inhibit the growth of the bacteria *in vitro* with a zone inhibition assay^45^. Sterile Whatman filters (Ø 5 mm) were placed on top of LB agar plates. These were then inoculated with 200 µL of permethrin at concentrations of 0, 10, 30 and 60 µM. We then pipetted 200 µL of a solution of 10^4^ CFU/mL of each bacterial strains on top of the filter, spreading through the agar. The plates were incubated at 27 °C (i.e., the temperature at which our experiments were performed). We detected no inhibition in any of the doses during the next 24 hours, suggesting that the four bacteria can tolerate the presence of permethrin *in vitro*.

To test if the bacterial treatments affect the permethrin-resistance of the two mosquito colonies we exposed larvae as described above and measured the resistance to permethrin five days after emergence with standard WHO tubes. For each treatment, five replicate groups of approximately 15 females were placed in tubes lined with 0.75% permethrin-impregnated paper (WHO standard paper) for 10 and 30 min for Kisumu and RSP colonies, respectively. We chose these times of exposure because preliminary experiments had shown that they would give about 50% mortality within 24 hours. Permethrin resistance was defined as the proportion of live mosquitoes per treatment 24h after exposure (Fig. 4).

### Statistical analysis

Statistical analyses were performed with the packages “DHARMa”^46^, “car”^47^, “survival”^48^, “lme4”^49^, and “emmeans”^50^ of R version 4.3.1 and figure panels assembled with Biorender.com.

A generalized linear model with a binomial error structure was to measure differences in individual developmental traits, for each of the mosquito colonies, due to bacterial treatment. To test for differences in infection persistence in alive individuals among bacterial treatments, we used a linear model with log-transformed bacterial load as the response variable, and treatment, time-point and their interaction as explanatory factors. For dead individuals, treatment was the only explanatory variable. Longevity across treatments for a given mosquito colony was compared with a cox proportional hazard with treatment as an explanatory factor. Insecticide resistance was analysed with a generalized linear mixed model with a binomial distribution of the errors for each of the mosquito colonies with alive/dead as the response variable. Treatment was the explanatory variable, while the replicate tube was considered as a random factor. Post-hoc multiple comparisons were performed with “emmeans” with the default Tukey adjustment.

## Acknowledgements

We thank Tiago G. Zeferino for his advice, technical support, and comments on the earlier drafts. LMS and the project were supported by SNF grant 310030_192786.

## Author contributions

LMS conceived the idea. JCK and LMS designed the experiments, and GA, LMS and MA performed them. LMS analysed the data and wrote the first draft of the manuscript. All authors contributed to the drafts.

## Data availability

The datasets and code generated in this study are available as Supplementary information files.

## Supplementary information

**Table S1.** Developmental traits.

| **Tested effect** |  |  |  |
| --- | --- | --- | --- |
| **Model 1a - Larvae mortality Kisumu** | **df** | **$\chi^2$** | ***p*** |
| Bacterial treatment | 4 | 7.833 | 0.098 |
| **Model 1b - Larvae mortality RSP** | **df** | **$\chi^2$** | ***p*** |
| Bacterial treatment | 4 | 12.796 | **0.012** |
| **Model 1c - Pupae mortality Kisumu** | **df** | **$\chi^2$** | ***p*** |
| Bacterial treatment | 4 | 0.184 | 0.996 |
| **Model 1d - Pupae mortality RSP** | **df** | **$\chi^2$** | ***p*** |
| Bacterial treatment | 4 | 3.561 | 0.469 |
| **Model 1e - Sex ratio Kisumu** | **df** | **$\chi^2$** | ***p*** |
| Bacterial treatment | 4 | 1.350 | 0.853 |
| **Model 1f - Sex ratio RSP** | **df** | **$\chi^2$** | ***p*** |
| Bacterial treatment | 4 | 3.923 | 0.417 |

**Table S2.**
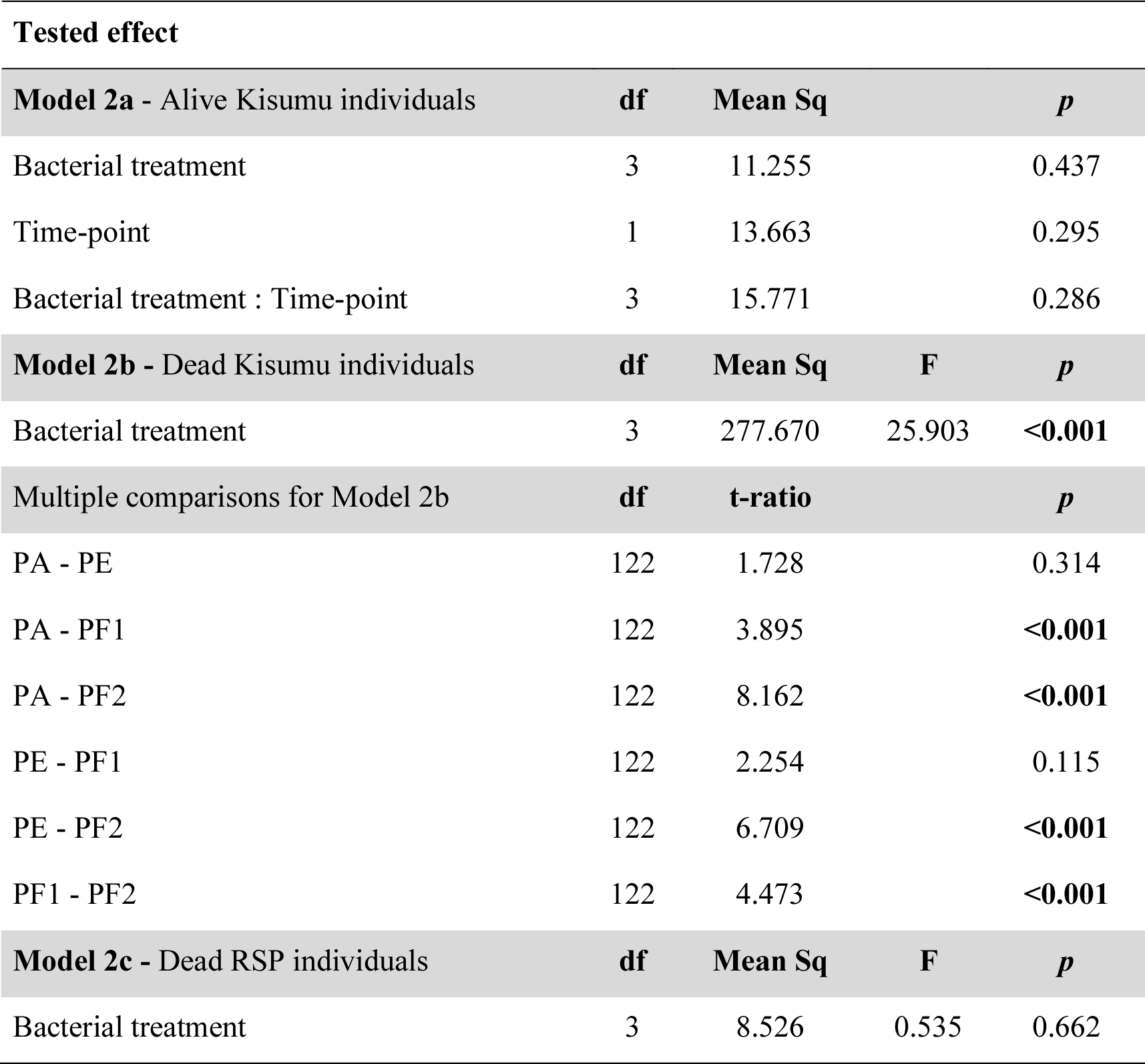
Bacterial persistence.

**Table S3.** Longevity.

| **Tested effect** |  |  |  |
| --- | --- | --- | --- |
| **Model 3a - Kisumu** | **df** | **$\chi^2$** | ***p*** |
| Bacterial treatment | 4 | 3.281 | 0.512 |
| **Model 3b - RSP** | **df** | **$\chi^2$** | ***p*** |
| Bacterial treatment | 4 | 3.049 | 0.550 |

**Table S4.** Insecticide resistance.

| **Tested effect** |  |  |  |
| --- | --- | --- | --- |
| **Model 4a - Kisumu** | **df** | **$\chi^2$** | ***p*** |
| Bacterial treatment | 4 | 19.847 | **< 0.001** |
| Multiple comparisons for Model 4a |  | **z-ratio** | ***p*** |
| Control - PA |  | 2.720 | 0.051 |
| Control - PE |  | 2.897 | **0.031** |
| Control - PF1 |  | 0.510 | 0.986 |
| Control - PF2 |  | -0.217 | 0.999 |
| PA - PE |  | 0.150 | 0.999 |
| PA - PF1 |  | -2.324 | 0.137 |
| PA - PF2 |  | -3.137 | **0.015** |
| PE - PF1 |  | -2.507 | 0.089 |
| PE - PF2 |  | -3.335 | **0.008** |
| PF1 - PF2 |  | -0.775 | 0.938 |
| **Model 4b - RSP** | **df** | **$\chi^2$** | ***p*** |
| Bacterial treatment | 4 | 24.802 | **< 0.001** |
| Multiple comparisons for Model 4b |  | **z-ratio** | ***p*** |
| Control - PA |  | 3.054 | **0.019** |
| Control - PE |  | 4.186 | **<0.001** |
| Control - PF1 |  | 4.263 | **<0.001** |
| Control - PF2 |  | 2.526 | 0.085 |
| PA - PE |  | 1.121 | 0.796 |
| PA - PF1 |  | 1.313 | 0.683 |
| PA - PF2 |  | -0.678 | 0.961 |
| PE - PF1 |  | 0.233 | 0.999 |
| PE - PF2 |  | -1.855 | 0.342 |
| PF1 - PF2 |  | -2.024 | 0.254 |

**Figure S1.**
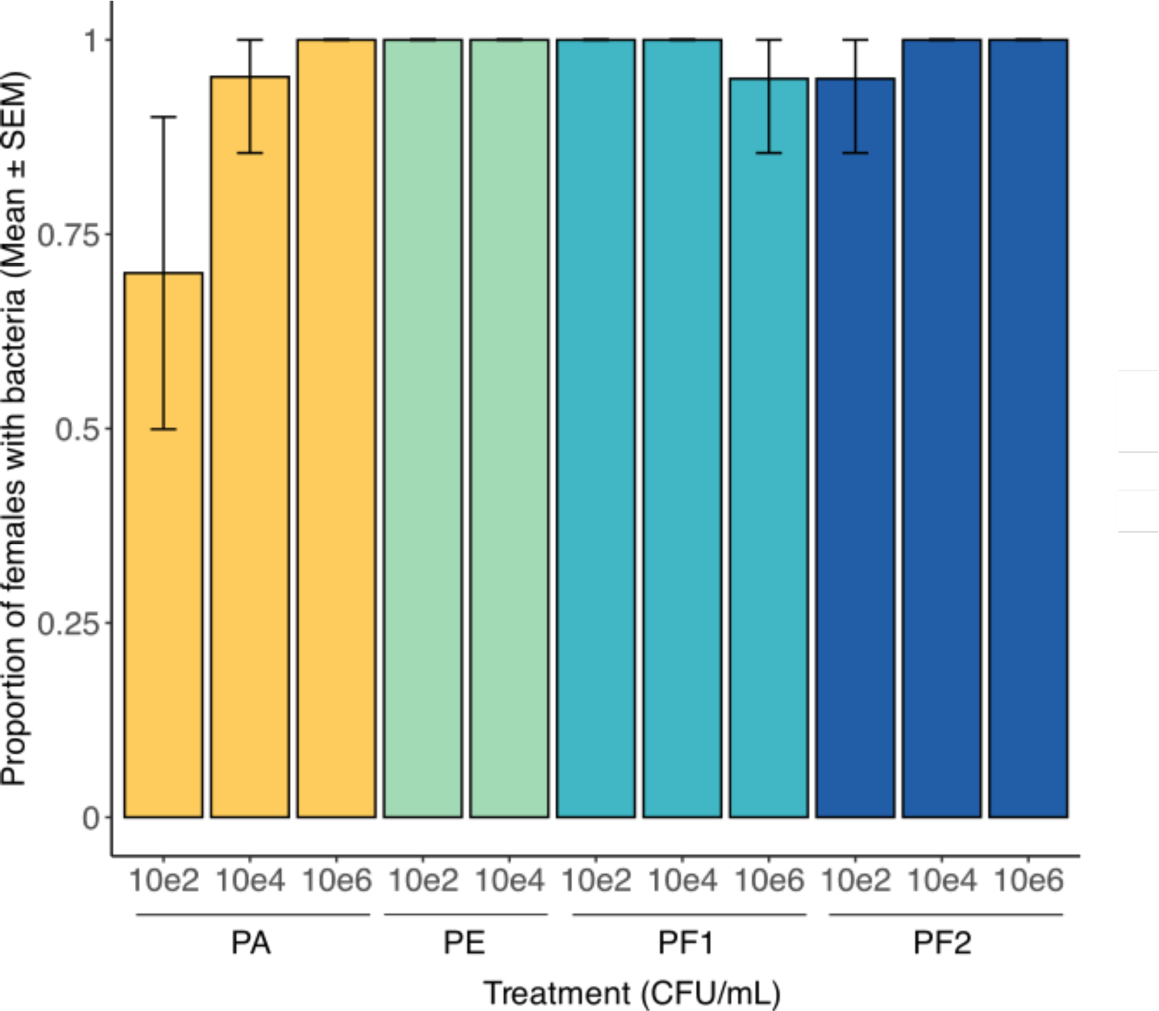
Infection success at different doses. Individuals were reared in distilled water with one of four bacterial strains, or an uninfected control, at one of the three doses (i.e., 10^2^, 10^4^ or 10^6^ CFU/mL). Bacterial load was assessed at day three post adult emergence on 20 Kisumu females per treatment and dose to confirm the bacteria would successfully colonize and grow through mosquito pupation. All the treatments survived to day three except for individuals infected with *P. entomophila* at the dose of 10^4^ CFU/ml, which died during the larval stage. A generalized linear model with a binomial error structure was used to test the effect of bacterial strain and dose on the likelihood of being infected. An interaction between both factors was also considered. Both an effect of treatment (χ2 = 13.153, *df* = 3, *p* = 0.004) and dose (χ2 = 8.3357, *df* =2, *p* = 0.015) were detected but no interaction between them (χ2 = 6.879, *df* = 5, *p* = 0.230).

